# A Novel Insertion Site for Group I Introns in tRNA Genes of Patescibacteria

**DOI:** 10.1101/2025.07.03.662931

**Authors:** Yuna Nakagawa, Kazuaki Amikura, Kimiho Omae, Shino Suzuki

**Author notes:** Address correspondence to Shino Suzuki.

## Abstract

Patescibacteria is a bacterial phylum with small genomes, frequent loss of essential genes, and the presence of introns. While many aspects of Patescibacteria remain enigmatic, an intriguing feature is the widespread occurrence of introns within their compact genomes. To better understand the diversity, roles, and evolution of bacterial introns, we focused on tRNA introns and analyzed Patescibacteria complete genome. Notably, 20% of these genomes lacked at least one tRNA gene for a canonical amino acid, primarily tRNA^Asn^ and tRNA^Asp^, whereas other tRNA genes were readily detected. This observation led us to conduct further analyses, resulting in the discovery of a novel group I intron insertion site at position 35/36 within the anticodon loop that likely prevented detection by conventional annotation tools. Splicing assays demonstrated that these bacterial introns are catalytically active and capable of self-splicing. To assess the broader distribution of this insertion site across bacteria, we analyzed 4,934 bacterial genomes and identified 269 group I introns within tRNA genes across 14 phyla. Nearly 70% of introns at position 35/36 originate from Patescibacteria, indicating that this feature is largely confined to the phylum. Subgroup classification showed that 79% of all tRNA introns belonged to the IC subgroup, whereas almost all Patescibacteria introns were assigned to IA, suggesting a distinct evolutionary origin. As most tRNA introns lacked homing endonuclease genes, horizontal transfer appears limited. Collectively, these findings advance our understanding of the phylogenetic distribution and evolutionary history of bacterial group I introns in tRNAs, with particular emphasis on Patescibacteria.

**IMPORTANCE:** Group I introns in bacterial tRNA genes were previously known only in a limited number of phyla. Our study expands this knowledge by identifying a novel insertion position in tRNA genes of phylum Patescibacteria and mapping their phylogenetic distribution across bacterial lineages. Our result revealed that group I introns inserted in tRNA genes differed in subgroups between Patescibacteria and other bacteria, highlighting the evolutionary uniqueness of introns of Patescibacteria. Additionally, we found that group I introns are maintained in 43% of bacterial phyla, with tRNA insertions being the most common. Our findings highlight that even in complete genomes, the presence of group I introns can hinder the detection of all 20 canonical tRNA genes by conventional tRNA annotation tools. This study illustrates the overlooked phylogenetic distribution of group I introns across the bacterial domain.

## INTRODUCTION

Recent advances in genome-resolved metagenomics have uncovered numerous previously unrecognized microbial lineages, fundamentally reshaping our understanding of microbial diversity and evolution. Among these, the phylum Patescibacteria, previously referred to as the Candidate Phyla Radiation (CPR), has drawn particular attention due to its remarkable diversity (1), uncertain phylogenetic position within the bacterial domain (2) and unusual biological features (3). Patescibacteria are a diverse clade of largely uncultivated organisms that are widely distributed across diverse environments, including groundwater aquifers (4), soils (5), freshwater systems (6), and oral microbiomes (7). Patescibacteria are characterized by ultra-small cell sizes (approximately 0.2–0.3 µm) and significantly reduced genomes ranging from 0.5 to 1.5 Mbp (8–10). Genomic analyses indicate that these organisms lack essential biosynthetic pathways and rely on interactions with other microbes for survival, with the few members that have been cultivated showed that they were epibionts on the surface of host microbes (7, 11–14). In addition to their metabolic streamlining, Patescibacteria exhibit atypical ribosomal features, including l including the lack of a subset of ribosomal proteins (3, 15), possessing frequent introns (1, 16) and encoding reduced sets of biogenesis factors (17). Given these atypical characteristics, Patescibacteria are considered a trove of unexplored biology, in terms of physiology, immune systems, evolutionary history, and translational mechanisms.

While many aspects of Patescibacteria remain enigmatic, one intriguing feature is the widespread presence of introns within their compact genomes. Group I and group II introns are self-splicing ribozymes, catalytically active RNA elements capable of excising themselves from precursor transcripts without the need for protein cofactors. In most bacterial species, rRNA and tRNA genes are typically compact and intronless, with only rare exceptions (3, 10, 18–22). However, members of the phylum Patescibacteria, which belong to the domain Bacteria, frequently harbor self-splicing introns within rRNA genes (3, 16) and, more rarely, within tRNA genes (10). The retention of these introns in the highly reduced genomes of Patescibacteria suggests that, despite the evolutionary pressure to minimize nonessential sequences, these elements may play important functional or regulatory roles in gene expression or genome evolution. Furthermore, group I introns are known to possess mobility, which may further contribute to genome dynamics through intron gain and loss events over evolutionary timescales (23–25).

Here, to further gain insight into the evolutionary and functional significance of intron-containing tRNAs, we thoroughly analyzed group I introns in tRNAs of domain Bacteria using complete genomes from the Genomic Taxonomy Database (GTDB) release 220 (26) that must encode at least 20 tRNAs. Our analysis revealed a diversity of group I introns in tRNAs. Notably, some tRNA^Asn^ and tRNA^Asp^ genes initially appeared to be absent but were actually present and disrupted by introns inserted at novel sites within the anticodon loop. In vitro assays confirmed that these introns are catalytically active and capable of producing mature tRNAs via self-splicing. Comprehensive genomic analysis of group I introns in tRNAs demonstrates that even the highly reduced genomes of Patescibacteria can retain functional self-splicing tRNA introns, which may play important roles in gene regulation and genome evolution or persist as relics of ancient genomic architecture.

## RESULTS AND DISCUSSION

### Some tRNA^Asn^ and tRNA^Asp^ Genes in Patescibacteria Are Not detected by Annotation Tools

We analyzed 95 Patescibacteria genomes, all classified as “complete genomes” in GTDB r220 (genome information listed in Supplementary Table 1), with each genome comprising a single contig. To assess the presence of tRNA genes corresponding to all 20 canonical amino acids, we employed tRNAscan-SE 2.0 (27). Our analysis revealed that 20.0% of these genomes did not encode a full set of tRNA genes covering all 20 canonical amino acids (Supplementary Table 2). Notably, tRNA^Asn^ genes were absent in 13.7% of the genomes, and tRNA^Asp^ genes were absent in 5.3% (Fig. 1a). We further validated with ARAGORN (28), a tool for detecting tRNA and tmRNA genes in nucleotide sequences. ARAGORN similarly failed to detect tRNA^Asn^ genes in 9.5% and tRNA^Asp^ genes in 4.2% of genomes (Fig. 1a, Supplementary Table 2). These absences were observed across multiple classes within the phylum Patescibacteria.

**Fig. 1.**
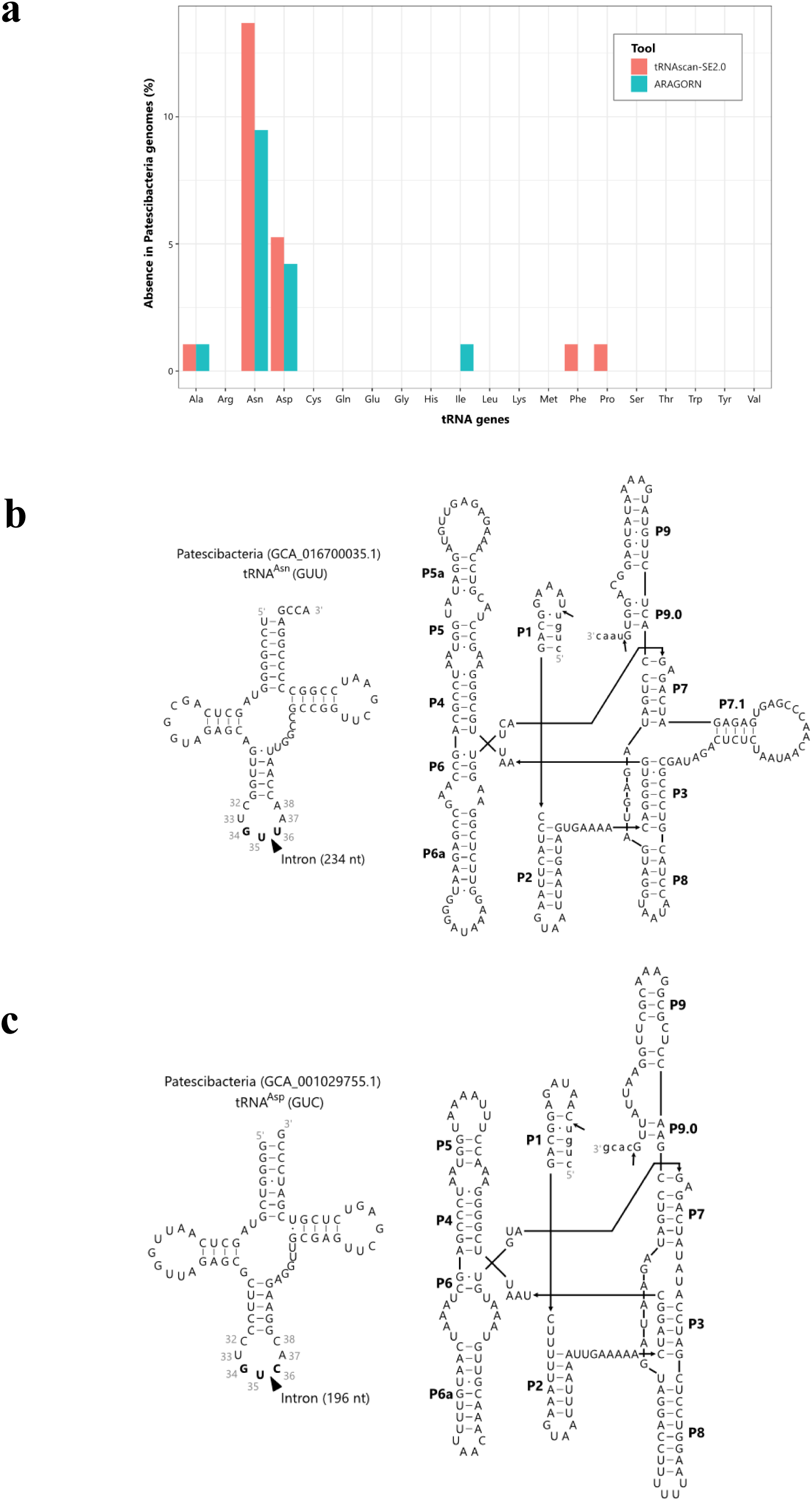
Identification of group I introns inserted at position 35/36 in tRNA^Asn^ and tRNA^Asp^ genes of Patescibacteria. (a) Percentage of Patescibacteria genomes in which specific tRNA genes were not detected using tRNAscan-SE2.0 (red) and ARAGORN (blue). The analysis includes all 95 genomes of Patescibacteria registered as complete genomes in GTDB r220. The x-axis shows tRNA genes corresponding to the 20 canonical amino acids, and the y-axis shows the percentage of genomes lacking each tRNA gene. (b) Predicted secondary structures of the tRNA^Asn^ (left) and the group I intron inserted into it (right) from the Patescibacteria genome (accession: GCA_016700035.1). (c) Predicted secondary structures of the tRNA^Asp^ (left) and the group I intron inserted into it (right) from the Patescibacteria genome (accession: GCA_001029755.1).

### Group I Intron Insertions at the Position 35/36 in tRNA^Asn^ and tRNA^Asp^

Given that complete genomes are expected to encode all the tRNA genes corresponding to the 20 canonical amino acids, we further investigated four tRNA^Asn^ genes and one tRNA^Asp^ gene that were not detected by tRNAscan-SE 2.0 but were identified by ARAGORN. Upon closer examination, we found that these genes contain introns. Using Infernal in conjunction with covariance models from the Rfam database (29), we determined that these introns belong to group I. These findings led us to hypothesize that group I introns may have prevented the detection of tRNA^Asn^ and tRNA^Asp^ genes in the 12 genomes where both tRNAscan-SE 2.0 and ARAGORN failed to identify them. Using the covariance model (RF00028.cm) for group I introns, we applied cmsearch from the Infernal package (30) (e-value < 1E−4) and detected putative group I introns in 11 out of the 12 genomes (Supplementary Table 3). To evaluate whether these introns were associated with tRNA gene fragments, we conducted BLAST searches (31) targeting regions flanking identified introns. In 10 out of the 11 genomes, the group I introns were found adjacent to partial tRNA^Asn^ and tRNA^Asp^ genes.

To determine the precise insertion site, we manually curated the intron boundaries by examining conserved structural features of group I introns. Specifically, we identified the conserved uridine residue at the end of the 5′ exon as a key marker guided by the well-characterized U–G base pairing that typically forms between the 5′ exon and the internal guide sequence in group I introns (32). Additionally, we identified a guanine residue at the 3′ end of the intron, which is another canonical feature of group I introns (33), as a complementary indicator to refine intron boundary determination. This analysis confirmed that in all 10 genomes, the introns were inserted at position 35/36 within the anticodon loop of tRNA^Asn^ (GUU) or tRNA^Asp^ (GUC) genes (Fig. 1b, 1c, Supplementary Table 4).

Castelle et al. (10) reported that introns in tRNA genes of several Patescibacteria were classified into three types. The first type includes canonical group I introns found in *Ca.* Microgenomates, *Ca.* Parcubacteria, and *Ca.* Peregrinibacteria, typically inserted one base after the anticodon (Position 37/38). The second type, called “divergent group I,” includes 15 introns similar in size and structure to group I but not detected by Rfam. These occur only in threonine tRNAs with the GGT anticodon, end in guanine, and are predicted to splice directly after the anticodon. The third type consists of small introns with defined secondary structures. However, because positions 35 and 36 within the anticodon loop of the tRNA^Asn^ (GUU) and tRNA^Asp^ (GUC) genes were not reported in the previous study, we decided to confirm them experimentally.

### Experimental Validation of Splicing Activity of Group I Introns Inserted at Position 35/36

Since group I introns at position 35/36 in the anticodon loop have not been previously reported, we conducted in vitro splicing assays to determine whether group I introns predicted in silico are indeed inserted at this position and capable of functional self-splicing. We selected sequences of tRNA^Asn^ from Patescibacteria (GCA_016700035.1) genome and tRNA^Asp^ from Patescibacteria (GCA_001029755.1) genome (sequence details are provided in Supplementary Table 5) as representatives. The GCA_016700035.1 genome encodes no intron-less tRNA^Asn^ genes and GCA_001029755.1 genome encodes no intron-less tRNA^Asp^ genes, which is consistent with other genomes predicted to harbor introns at position 35/36 in the anticodon loop of tRNA^Asn^ or tRNA^Asp^ genes. Thus, in both cases, the intron-containing tRNA genes represent the sole genomic copies of these tRNA genes, indicating that splicing is essential to produce functional tRNAs. As a positive control in the in vitro splicing assay, we used a tRNA^Leu^ (UAA) from a genome of Cyanobacteriota (GenBank ID: BA000019.2) which contains a group I intron previously demonstrated to splice efficiently under in vitro conditions (19). Upon successful splicing, distinct bands corresponding to splicing intermediates (IVS-3’ exon), spliced tRNA (ligated exon), linear intron (L-IVS), and circular intron (C-IVS) are expected to appear (19) (Fig. 2a). In the positive control, these bands were clearly observed, confirming the reliability of our experimental system (Fig. 2b).

**Fig. 2.**
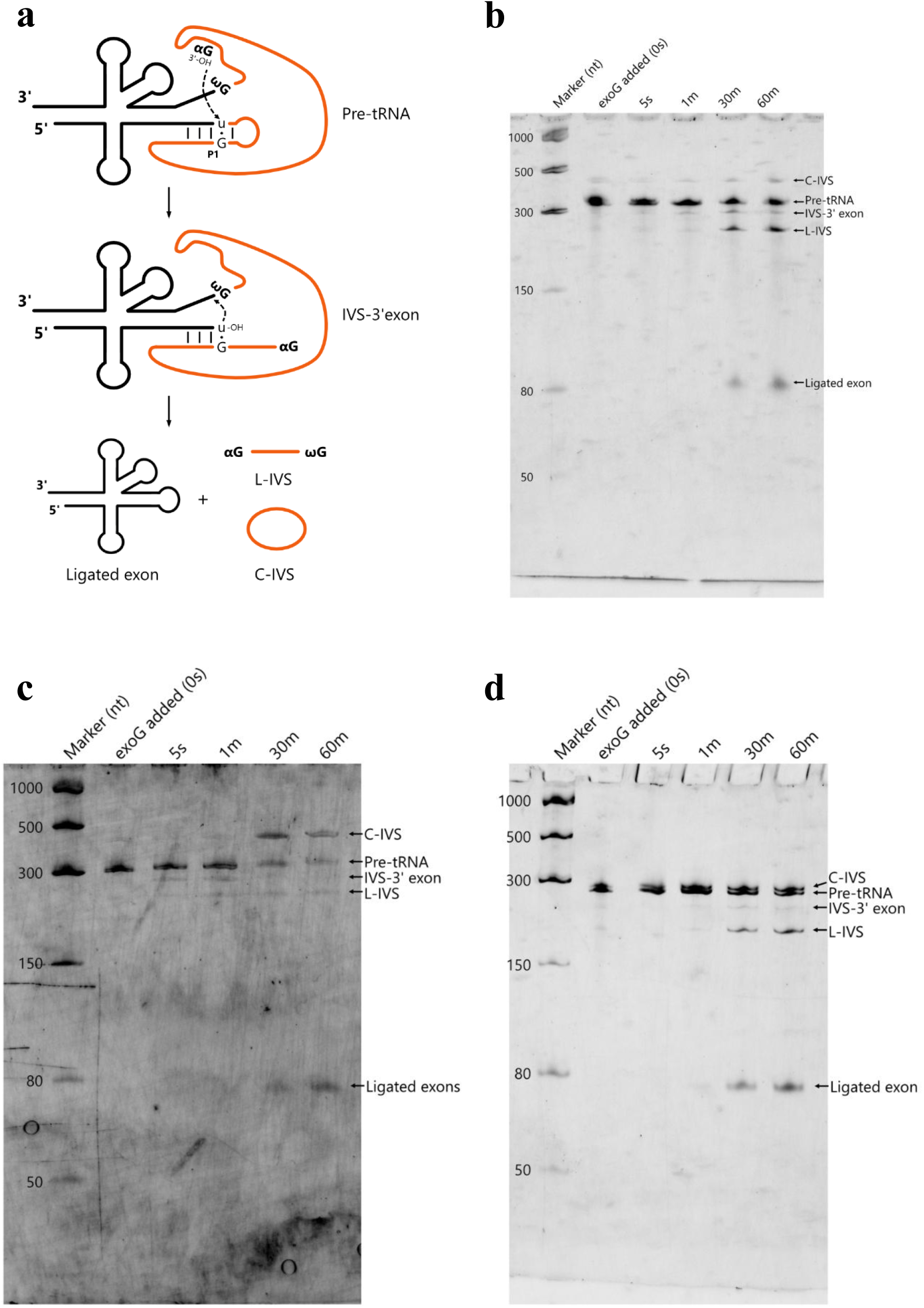
Splicing activity of group I introns inserted at position 35/36 in tRNA genes of Patescibacteria. (a) Schematic drawing of the splicing reactions of group I introns in tRNA genes. ω G: A 3’-terminal guanosine of the intron. α G: An exogenous guanosine cofactor. L-IVS: A linear intron. C-IVS: A circular intron. (b) Gel electrophoresis of splicing reactions of the tRNA^Leu^ containing a group I intron from the Cyanobacteriota genome (GenBank ID: BA000019.2). (c) Gel electrophoresis of splicing reactions of the tRNA^Asn^ containing a group I intron from the Patescibacteria genome (accession: GCA_016700035.1). (d) Gel electrophoresis of splicing reactions of the tRNA^Asp^ containing a group I intron from the Patescibacteria genome (accession: GCA_001029755.1). (b-d) Time points after GTP addition are indicated above each lane (0 seconds, 5 seconds, 1 minute, 30 minutes and 1 hour).

In the constructs of tRNA^Asn^ and tRNA^Asp^ from Patescibacteria genomes, distinct bands corresponding to spliced products were observed after 30 minutes of incubation (Fig. 2c, d). To confirm the splicing junctions, the ligated exons were extracted and sequenced, revealing mature tRNA^Asn^ and tRNA^Asp^ sequences, confirming that the introns were accurately excised at position 35/36 within the anticodon loop. These results were the first to experimentally demonstrate the splicing activity of group I introns both at a previously unreported insertion site in bacteria and within the genomes of Patescibacteria.

### Insertions at position 35/36 Are Unique in a phylum Patescibacteria

To determine whether group I introns at position 35/36 in Patescibacteria are widely distributed across other bacterial phyla, we expanded our analysis to extend all bacterial genomes. Using the same approach that had successfully identified group I introns in tRNA genes of Patescibacteria, we examined all 4,934 bacterial genomes (spanning 63 phyla) classified as complete genomes in GTDB r220. This comprehensive analysis identified 269 group I introns inserted into tRNA genes across the bacterial genomes (Supplementary Table 4).

In Patescibacteria, we observed multiple genomes in which only intron-containing tRNA^Asn^ and tRNA^Asp^ genes were detected, with no corresponding intron-less copies. A similar trend was found across the bacterial domain, where approximately 90% of the 269 group I intron-containing tRNA genes identified lacked an intron-less counterpart with the same anticodon. Thus, it is rare for both intron-containing and intron-less copies of the same tRNA gene to coexist within a single genome. Therefore, when complete genomes lack one or more of the canonical tRNA genes, we should consider the possibility of intron insertions.

We then analyzed the insertion positions of these group I introns within tRNA genes. In agreement with previous studies (18–20), insertions were commonly found at positions such as 33/34 in tRNA^fMet^, 34/35 in tRNA^Leu^, and 36/37 in tRNA^Arg^ (Fig. 3a). Beyond these well-characterized sites, we found that group I introns were inserted at nearly all positions within the anticodon loop except for position 37/38. Among the group I introns newly identified in this study at position 35/36, a total of 28 introns were found at this position across all bacterial genomes.

**Fig. 3.**
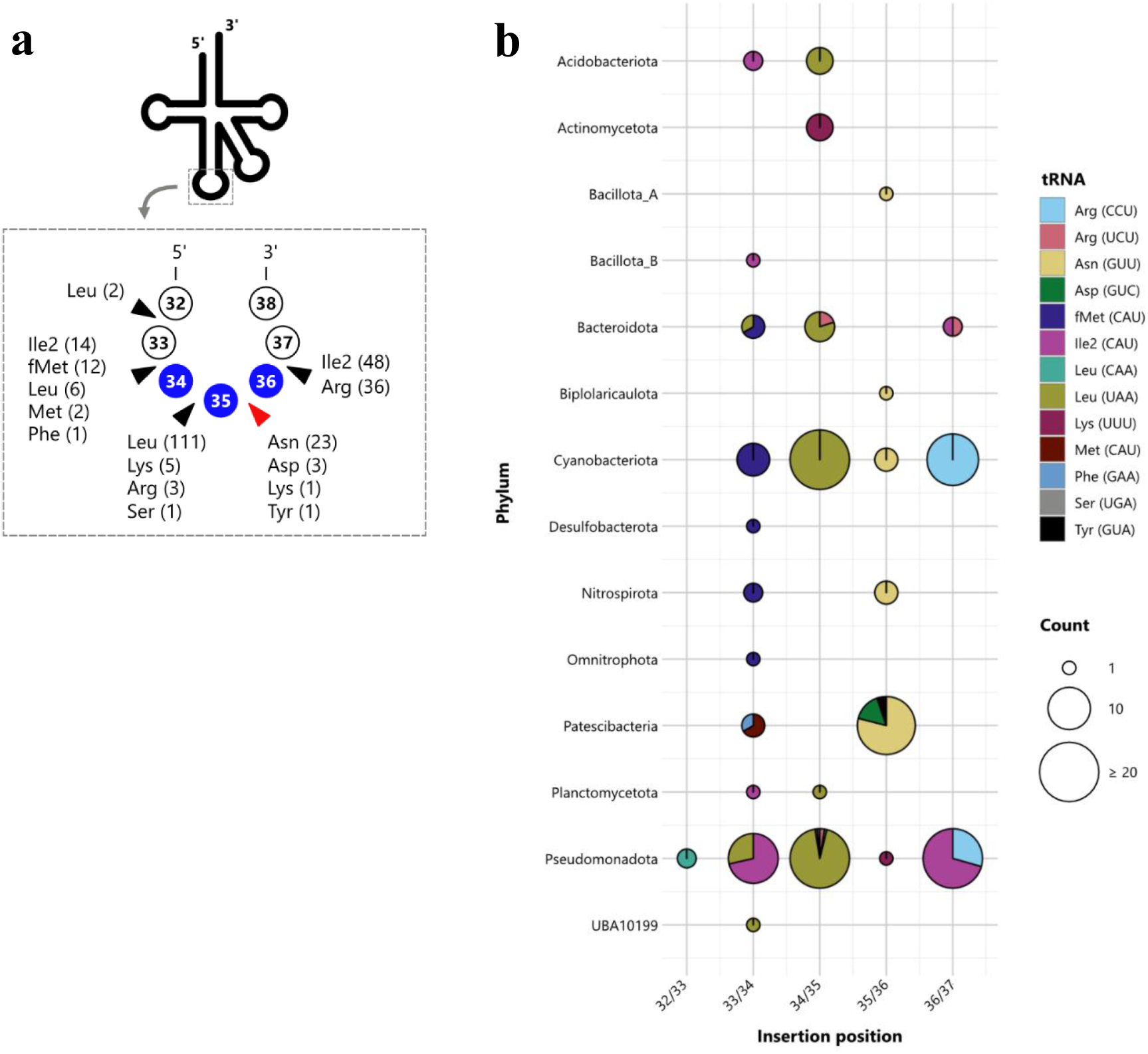
Diversity of insertion sites of group I introns in bacterial tRNA genes. (a) Schematic representation showing the various insertion sites of group I introns within the anticodon loop of tRNA genes. The newly identified position 35/36 discovered in this study is indicated by a red triangle. Numbers in parentheses represent the number of group I introns identified at each position for each amino acid–specific tRNA gene. (b) Bubble chart showing the insertion positions of group I introns within tRNA genes across bacterial phyla. Each circle represents the number of genomes with introns inserted at a specific position (x-axis) and in a particular phylum (y-axis). Circle size corresponds to the number of observations (legend at bottom right). Colors indicate the amino acid and anticodon of the associated tRNA genes.

When we examined the distribution of insertion positions by a phylum, we found that in the phylum Pseudomonadota, which had the highest number of tRNAs containing group I introns, and in the phylum Cyanobacteriota, which had the second highest, the introns were primarily inserted at positions 34/35 and 36/37 (Fig. 3b). In contrast, Patescibacteria showed no insertions at these common positions, but instead exhibited a marked enrichment at the position 35/36. Of 28 tRNA introns inserted at the position 35/36, 19 were derived from Patescibacteria, while the remaining cases were found in the phyla Cyanobacteriota and Nitrospirota (3 each), and in Baccilota_A, Bipolaricaulota, and Pseudomonadota (1 each). These findings indicate that intron insertions at the position 35/36 were particularly enriched in Patescibacteria, distinguishing Patescibacteria from other bacterial phyla.

A previous study (10) identified tRNAs containing group I introns in multiple genomes of Patescibacteria using the eukaryotic settings of tRNAscan-SE. These introns were typically annotated as being inserted between positions 37 and 38, which corresponds to a common intron insertion site in eukaryotes (34). In our genomic analysis, in which the boundaries of tRNA-inserted group I introns were manually defined based on conserved structural information, we did not identify any introns at the position 37/38, suggesting that the previously reported annotations at this site were likely represented annotations introduced by the use of eukaryotic prediction settings. When we input some tRNA sequences with introns inserted at positions other than 37/38 into tRNAscan-SE 2.0 using the eukaryotic settings, the intron was consistently predicted to be at position 37/38. This result indicates that while the presence of introns can be detected, accurate determination of the insertion site for group I introns requires additional manual adjustment and validation based on structural information with the current bioinformatic tool.

### tRNA Introns of Patescibacteria Encode Homing Endonucleases of HNH family

We investigated whether homing endonuclease genes (HEGs), which are often associated with horizontal gene transfer, are encoded in the group I introns inserted into tRNA genes. Group I introns longer than 500 bp are known to potentially encode HEGs (35). Therefore, we analyzed the length distribution of the identified group I introns and found that their lengths ranged from 184 to 894 base pairs, with a median length of 255 bp (Fig. 4a). Only 10 of the 269 introns exceeded 500 bp, and all these encoded HEGs (Supplementary Table 4). Group I introns within tRNA genes decoding the same anticodon tended to encode endonucleases from the same family. For example, three introns at position 36/37 in tRNA^Arg^ (CCU) from Cyanobacteriota encoded LAGLIDADG-type endonucleases; two introns at position 33/34 in tRNA^fMet^ (CAU) from Cyanobacteriota encoded PD-(D/E)XK domain-containing endonucleases; and one intron at position 34/35 in tRNA^Leu^ (UAA) from Bacteroidota and four introns at position 35/36 in tRNA^Asn^ (GUU) from Patescibacteria encoded endonucleases of the HNH family (Fig. 4a). Notably, the HEGs found in rRNA introns of Patescibacteria belong to the LAGLIDADG family (16), suggesting that tRNA and rRNA introns may have been acquired through distinct evolutionary routes. Importantly, most group I introns inserted into tRNAs did not encode HEGs. Since HEGs are often required for mobility and self-propagation of introns, their absence implies that these tRNA introns may lack autonomous mobility. This finding suggests that, at present, horizontal transfer of tRNA introns occurs only infrequently.

**Fig. 4.**
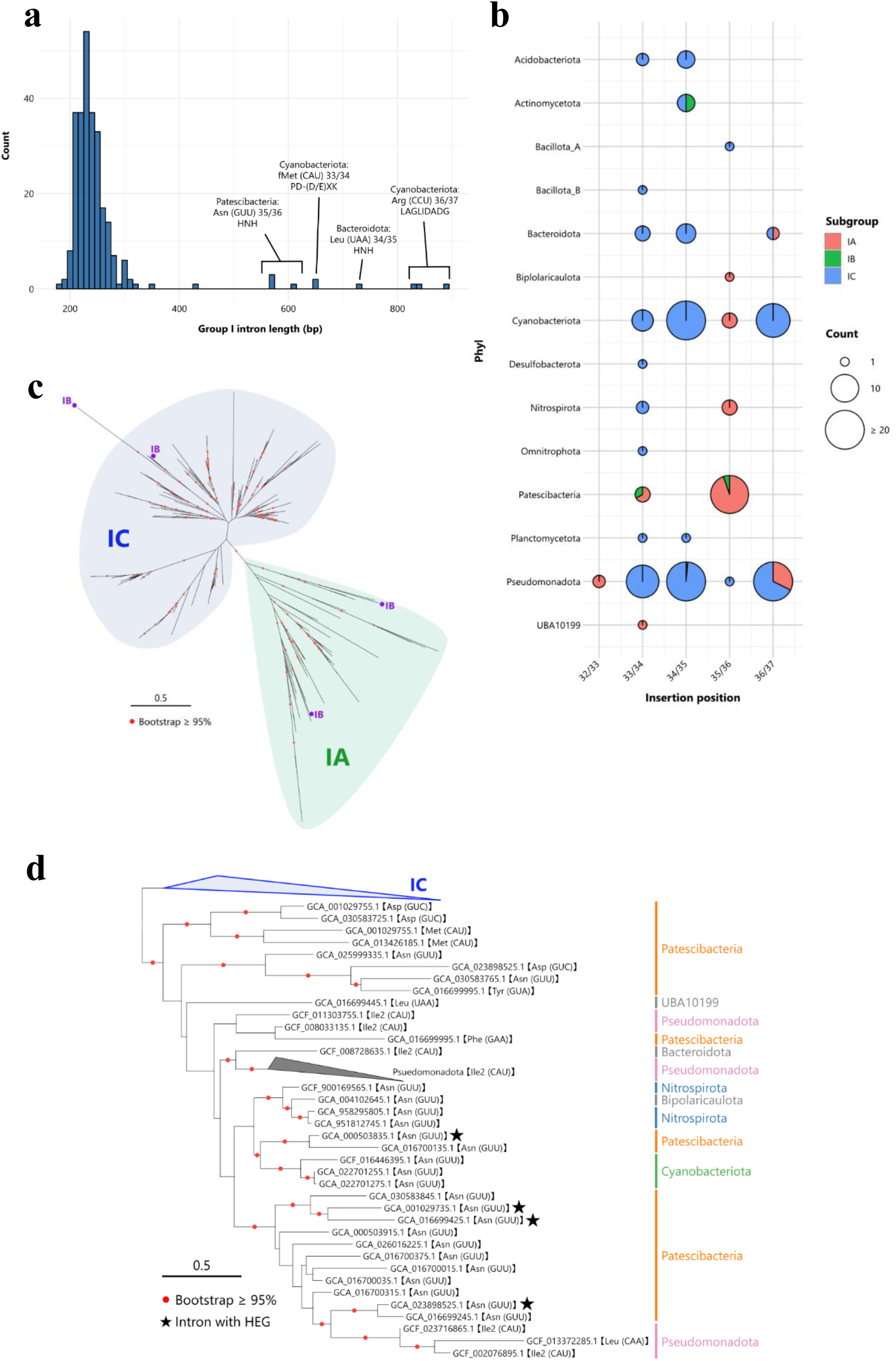
Distribution of subgroups of group I introns across insertion positions and bacterial phyla. (a) Histogram showing the length distribution of group I introns identified from bacterial tRNA genes. Introns longer than 500 bp contained HEGs. Among these, HNH-family HEGs were identified in the tRNA^Asn^ (GUU) of Patescibacteria and the tRNA^Leu^ (UAA) of Bacteroidota; the PD-(D/E)XK family was found in the tRNA^fMet^ (CAU) of Cyanobacteriota; and the LAGLIDADG family was observed in the tRNA^Arg^ (CCU) of Cyanobacteriota. (b) Bubble chart illustrating the distribution of subgroups of group I introns inserted at different positions within tRNA genes across bacterial phyla. Each circle represents the number of introns at a given insertion site (x-axis) in a particular phylum (y-axis), with the size of the circle indicating the count. The color within each circle denotes the proportion of subgroups: IA (red), IB (green), and IC (blue). (c) Maximum-likelihood phylogenetic tree of 269 group I intron sequences inserted into bacterial tRNA genes. The tree was inferred using IQ-TREE with the TNe+R7 model selected by ModelFinder, based on 186 nucleotide sites. The tree shows a clear separation between the IA and IC subgroups, with bootstrap values ≥ 95% indicated by red dots. (d) Enlarged view of the IA clade from panel (c). Genome IDs and corresponding tRNA genes are labeled at the tips of the phylogenetic tree, and phylum-level annotations are provided for each branch. Stars indicate group I introns that contain HEGs.

### Group I Introns of Patescibacteria Represent a Distinct Structural Subtype

To investigate whether group I introns in Patescibacteria exhibit characteristics distinct from those found in other bacterial phyla, we analyzed the structural features of these introns. We classified the structural subgroups of each intron using Infernal with covariance models representing group I intron subgroups (35). Out of the 269 tRNA introns identified across bacteria, 78.8% belonged to the IC subgroup. However, 25 out of 28 introns inserted at the position 35/36 were classified into the IA subgroup (Fig. 4b). None of the group I introns from tRNA genes of Patescibacteria were assigned to the IC subgroup. This contrast suggests that Patescibacteria may have acquired their introns through an evolutionary trajectory that is distinct from those observed in other bacterial phyla. A comprehensive analysis of group I introns across all domains of life has shown that IC introns are the most abundant (35, 36). Within the IC subgroup, most commonly identified introns are found in plastid tRNA^Leu^ genes (36), which share an evolutionary relationship with Cyanobacteriota tRNA^Leu^ genes (20, 37). In comparison, IA introns are relatively rare and have mainly been reported in certain bacteria, protists (such as centrohelids and cryptophytes), fungi, and viruses (35). However, the distribution of IA introns across different classes of Patescibacteria suggests that these introns may have been acquired at an early stage following the divergence of Patescibacteria.

To further investigate the evolutionary relationships among these introns, we constructed a maximum-likelihood phylogenetic tree based on all tRNA intron sequences identified across bacteria. This analysis revealed that the introns clustered clearly into distinct subgroups, with the IA and IC classes forming well-supported clades (Fig. 4c). Within the IA clade, Group I introns inserted into tRNA genes corresponding to the same canonical amino acids tended to cluster together. For example, tRNA^Asn^ introns from Patescibacteria and Cyanobacteriota clustered within the same clade, while tRNA^Ile2^ introns from Bacteroidota and Pseudomonadota also formed a shared clade (Fig. 4d). Furthermore, introns from the same phylum often formed closely related clades. For instance, in the branches containing tRNA^Asn^ introns from both Patescibacteria and Cyanobacteriota, the two groups formed distinct subclusters. Interestingly, the tRNA^Asn^ introns from Patescibacteria also formed a sister clade with tRNA^Ile2^ and tRNA^Leu^ introns from Pseudomonadota, supported by high bootstrap values. Among the four tRNA^Asn^ introns from Patescibacteria that encoded HEGs, two were closely related, one clustered with Cyanobacteriota, and another with Pseudomonadota. These patterns suggest the possibility of HGT between these groups. Based on our data, group I introns inserted at position 35/36 and classified in the IA subgroup that encode homing endonucleases were found only in Patescibacteria. This raises the possibility that these introns were horizontally transferred from Patescibacteria to other phyla, or alternatively, that they were horizontally acquired from outside Patescibacteria and subsequently lost their homing endonucleases in other lineages. In this study, our analysis was limited to complete genomes to obtain solid understanding of intron distribution; however, expanding the dataset to include draft genomes may reveal additional group I introns that encode homing endonucleases, potentially allowing for more comprehensive phylogenetic analyses.

### tRNA Genes Are the Most Frequent Targets of Group I Intron Insertions in Bacteria

Finally, to investigate how group I introns are distributed not only in tRNAs but across the entire bacterial domain, we conducted a comprehensive search for group I introns. We analyzed group I introns using Infernal (e-value <1E-4) across 4,934 complete genomes representing 63 bacterial phyla. As a result, a total of 682 group I introns were identified in 441 genomes across 27 phyla (Supplementary Table 3). On average, genomes containing group I introns harbored 1.5 copies (standard deviation = 1.5, median = 1), indicating that most genomes conserved only a single intron. Group I introns were particularly enriched in the phyla Patescibacteria and Cyanobacteriota, where over one-third of the analyzed genomes conserved at least one group I intron (Fig. 5a). As a comparison to group I introns, we also investigated the distribution of group II introns using the same method and identified a total of 5,662 group II introns. The average copy number per genome was 5.2 copies (standard deviation = 13.4, median = 2) (Supplementary Table 3), suggesting a broader distribution and higher copy number per genome compared to group I introns. While the phylum Cyanobacteriota, which frequently contained group I introns, also showed a high prevalence of group II introns, no group II introns were detected in Patescibacteria.

**Fig. 5.**
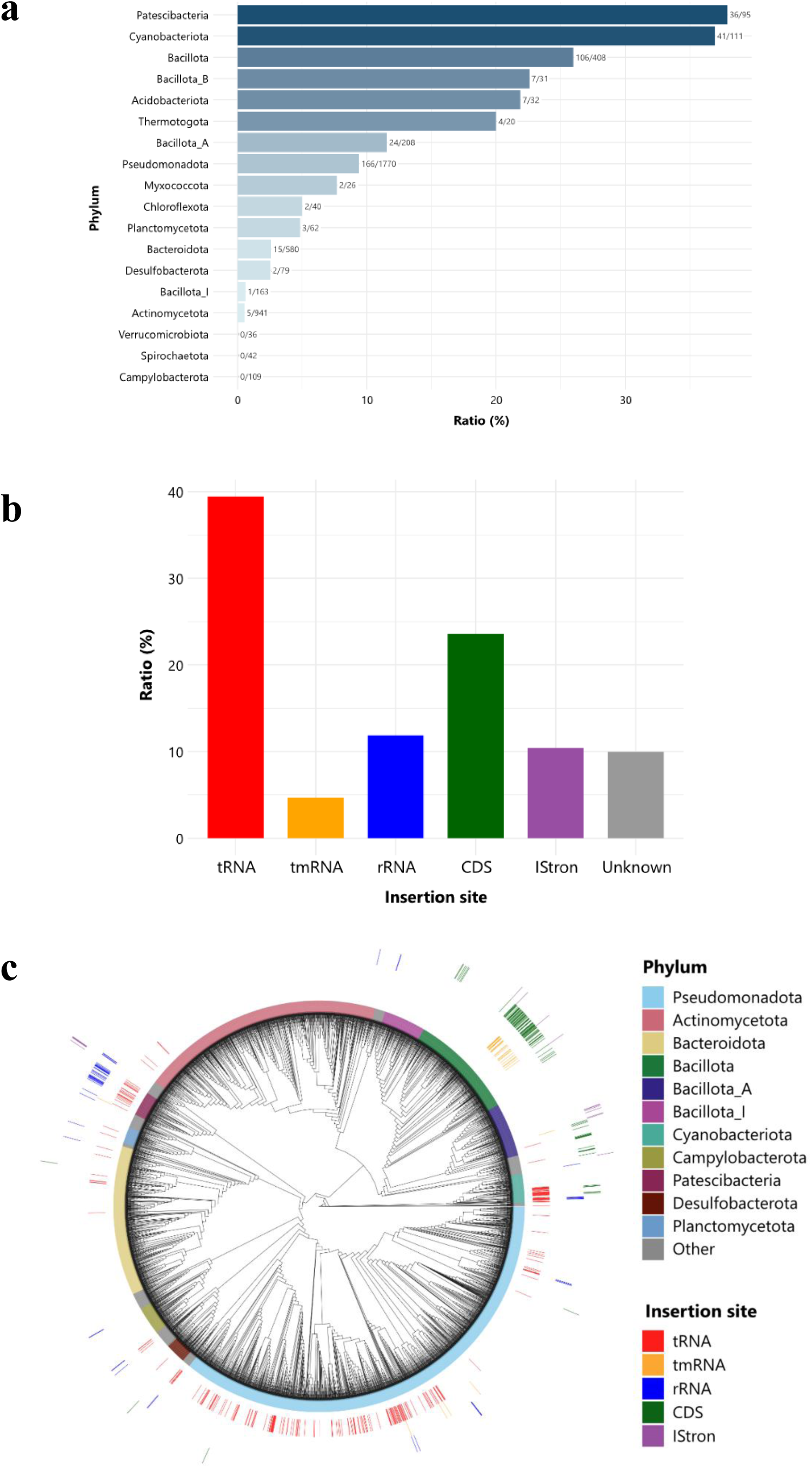
Distribution of insertion sites of group I introns across bacterial genomes. (a) Bar plot showing the proportion of genomes containing group I introns for each phylum. Only phyla with ≥20 genomes are included. The retention rate was calculated as the number of genomes with group I introns divided by the total number of genomes in each phylum. Numbers on the right side of each bar indicate the count of intron-containing genomes and the total number of genomes. (b) Bar graph showing the proportions of the 682 group I introns identified from bacterial genomes by their insertion sites: tRNA genes, tmRNA genes, rRNA genes, protein-coding sequences (CDSs), IStrons, and unknown loci. (c) Phylogenetic tree of 4,934 bacterial genomes constructed using concatenated alignments of bac120 marker genes. The tree was inferred using IQ-TREE with the Q.pfam+R10 model selected by ModelFinder, based on 5,035 amino acid positions. Phylum and insertion sites were mapped on the tree.

To determine the functional gene categories that harbor group I introns in bacterial genomes, we classified the genes that inserted all identified introns. As a result, tRNA genes were the most frequent targets of intron insertion, with 269 cases, which accounted for approximately 40% of all identified introns (Fig. 5b). 32 introns were inserted into tmRNA genes (Supplementary Table. 6), 81 into rRNA genes (Supplementary Table. 7), and 161 into protein-coding sequences (CDSs) (Supplementary Table. 8). In addition, 71 introns were identified near TnpB-like genes (Supplementary Table. 8). IStrons are known as genetic elements that combine a group I intron with a mobile insertion sequence (IS element), and have been reported to include TnpB genes, which encode RNA-guided DNA endonucleases (38, 39). Therefore, the introns found adjacent to TnpB-like genes were presumed to be associated with IStrons.

Group I introns in tRNA genes were identified across 14 of the 63 bacterial phyla analyzed, representing a phylogenetic diversity comparable to that of rRNA introns, which were found in 15 phyla (Fig. 5c). This finding suggests that introns are not limited to specific phyla but are rather broadly distributed across the bacterial domain. In Patescibacteria, group I introns are conserved across the entire phylum in both tRNA and rRNA genes, suggesting the presence of selective pressures or regulatory mechanisms that maintain these elements in this phylum. Although CDSs were the second most common insertion targets after tRNA genes, this was largely due to a few genomes harboring multiple intron-containing CDSs, most of which were found in the phylum Bacillota (Fig. 5c). Similarly, almost all tmRNA-associated introns were also observed in Bacillota (Fig. 5c). These results demonstrate that group I introns exhibit broad phylogenetic distribution in tRNA and rRNA genes, whereas their presence in CDS and tmRNA genes is more restricted.

Group I introns are generally known to predominately insert into structured RNAs such as rRNAs and tRNAs (25), and our findings are consistent with this tendency. This is likely because introns that insert into these evolutionarily conserved and stable genes tend to be retained in the genome (22) and the infrequent loss of introns may be related to the fact that the genes they insert into often exist as single copies. Notably, in both Patescibacteria and other bacteria, it was rare to find both intron-containing and intronless copies of tRNA genes encoding the same anticodon within a single genome. Since tRNA genes are essential for survival, mutations that disrupt either the intron or its splicing are likely to render the gene nonfunctional and therefore be subject to strong negative selection. Consequently, once an intron has been inserted into a tRNA gene, its loss is also likely to be disfavored by selection. Consistently, rRNA genes, which also frequently harbor group I introns, are typically present in only a single copy in Patescibacteria (Supplementary Table 9).

Currently, the retention rate of tRNA genes is widely used as an indicator of genome completeness in bacterial genome quality assessments (40). However, as demonstrated in this study, group I introns are retained in tRNA genes across multiple bacterial phyla, and their presence can interfere with the detection of these genes by the tRNA annotation tools, leading to underestimation of genome completeness. Therefore, future updates to bacterial tRNA annotation tools should aim to accommodate intron-containing tRNAs, including those with introns inserted at newly identified positions such as 35/36 within the anticodon loop.

The retention of seemingly dispensable genetic elements such as group I introns in Patescibacteria, which are characterized by highly reduced genomes, raises an intriguing question. It remains unclear whether Patescibacteria are particularly prone to acquiring group I introns or whether they are simply more likely to retain them once acquired. Interestingly, although both are self-splicing introns, group II introns have not been found in Patescibacteria, with only fragmented sequences occasionally detected (16). This suggests that only group I introns have successfully persisted in Patescibacteria. In contrast to Cyanobacteriota, which also retains group I introns but commonly harbors group II introns as well, this pattern implies that Patescibacteria may possess unique intron retention mechanisms or selection pressures specific to group I introns. Overall, our findings provide insights on the origins, evolutionary history, and adaptation strategies of Group I introns in bacteria, especially in Patescibacteria.

## MATERIALS AND METHODS

### Data set

4,934 representative bacterial genomes were downloaded from GTDB r220 (26) and filtered using the following criteria: (i) assembly level: complete genome; (ii) number of contigs: 1; and (iii) contamination: <5%.

### Identification of Group I / II Introns and rRNA genes

Group I introns, group II introns, and rRNA genes were detected using cmsearch from Infernal v1.1.4 (30) against covariance models from the Rfam database (29), with a threshold e-value of <1E-4. We used RF00028.cm for Group I introns, RF00029.cm for Group II introns, RF00177.cm for bacterial SSU rRNA, and RF02541.cm for bacterial LSU rRNA. The secondary structures of the identified group I introns were initially predicted using R2DT (41) and subsequently manually refined to improve clarity and visualization.

### Identification of Genes Containing Group I Introns

To identify genes disrupted by group I introns, we applied the following approaches:

i. tRNA genes: tRNA genes were predicted using tRNAscan-SE 2.0 with the -B option (27). For genomes of Patescibacteria, additional searches were conducted using ARAGORN v1.2.41 with the -i option (28) to detect potential intron-containing tRNAs. Intron-less tRNA sequences detected by tRNAscan-SE 2.0 were used to build a BLAST database with the makeblastdb command from BLAST+ v2.12. (31). Predicted intron regions from Infernal, including 100 bp upstream and downstream flanks, were used as queries. BLAST searches were performed using the blastn command with the -task blastn-short option. When tRNA sequences were detected on both sides of the intron, boundaries were manually curated.
ii. rRNA and tmRNA genes: Predicted introns with 1,000 bp flanking regions were queried against Rfam covariance models for bacterial SSU rRNA (RF00177.cm), bacterial LSU rRNA (RF02541.cm), and tmRNA (RF00023.cm) using cmsearch program in Infernal. To improve the detection of fragmented sequences, the --anytrunc option was used. For rRNA, when rRNA sequences were detected on both sides of a group I intron, the intron was considered to be inserted within the rRNA gene. For tmRNA, group I intron insertion caused the 3′ end of the tmRNA to become too short (42) to be detected by Infernal. Therefore, when only a partial tmRNA sequence was detected near one side of the intron, the boundary was manually inspected to confirm whether the intron was inserted within the tmRNA sequence.
iii. Protein-coding genes (CDS): The intron regions and their 4,000 bp flanking sequences were analyzed using the blastx command in DIAMOND (43) against the NCBI non-redundant protein database to investigate trends in genes frequently disrupted by intron insertions. Based on the initial results, we focused on recurrently observed genes previously reported in the literature (44–47), such as flagellin and ribonucleotide reductase, and downloaded representative sequences for these genes from InterPro (48). A custom database was then constructed, and additional blastx searches were performed using these sequences (see Supplementary Table 10 for details). When sequences were identified on both sides of an intron, or when a single flanking sequence was located immediately adjacent to the intron, they were considered as intron insertions within CDSs. For genes that were identified less frequently, we manually reviewed the annotations obtained from the initial blastx results.

### In Vitro Transcription and Splicing Assay

DNA templates for in vitro transcription were synthesized by Twist Bioscience and Eurofins (listed in Supplementary Table 5) and PCR-amplified using PrimeSTAR MAX DNA Polymerase (TaKaRa). For PCR amplification, a sequence containing the T7 promoter (5’-TAATACGACTCACTATA-3’) was added to the 5′ end of the forward primer, allowing the target sequence to be positioned immediately downstream of the promoter (see Supplementary Table 5 for details). For the tRNA^Asn^ construct from GCA_016700035.1, an additional guanine (G) was inserted between the T7 promoter and the tRNA sequence to enhance transcription initiation efficiency. PCR products were purified using the NucleoSpin Gel and PCR Clean-up kit (TaKaRa). In vitro transcription was performed using T7 RNA polymerase (New England Biolabs) following the manufacturer’s protocol. Following incubation at 37°C for 2 hours, 0.2 µl DNase I (Nippon Gene) was added and incubated for an additional 15 minutes at 37°C. Transcribed RNAs were purified using the RNA Clean & Concentrator kit (Zymo Research).

Precursor RNAs were resolved on a 7% UREA-PAGE gel (49) by electrophoresis at 180 V. A precursor band corresponding to a tRNA with intron was excised from the gel and incubated in 0.3 M NaCl with rotation for over 12 hours to elute the RNA (50). The eluted product was filtered and purified by ethanol precipitation.

Splicing assays were performed with reference to previous studies (19, 51). Prior to splicing, RNA samples were denatured at 95°C for 3 minutes. Samples were then incubated at 30°C in buffer (50 mM HEPES-KOH pH 7.5, 5 mM MgCl_2_, RNase-free water to 10 µl) containing 10–200 nM RNA. Splicing was initiated by the addition of 0.1 mM GTP and terminated at 5 seconds, 1 minute, 30 minutes, and 1 hour by adding 1 µl of 0.3 M EDTA.

### Validation of Splicing by RT-PCR and Sequencing

RNA products corresponding to the predicted mature tRNA band were excised and purified from the gel after an hour incubation in the splicing assay, following the same procedure described above. Reverse transcription was performed using PrimeScript Reverse Transcriptase (TaKaRa) according to the manufacturer’s protocol, with random primers. Prior to the reaction, the RNA templates were denatured at 95°C for 3 minutes.

To enable sequencing of the tRNA sequence (less than 80 bp) by Sanger sequencing, PCR amplification of the cDNA was performed (see Supplementary Table 5) with PrimeSTAR MAX DNA Polymerase (TaKaRa). The PCR products were then further amplified using the forward primer (5’-GCTATTTAGGTGACACTATAG-3’) and the reverse primer (5’-AATACGACTCACTATAGG-3’), and the resulting fragments were gel-purified using the NucleoSpin Gel and PCR Clean-up kit (TaKaRa). The purified products were confirmed by Sanger sequencing.

### Construction of Phylogenetic Tree for Bacterial Genomes

Phylogenetic analysis was conducted using bac120 marker gene alignments provided by GTDB r220. Maximum-likelihood phylogenetic trees were constructed using IQ-TREE v2.1.4-beta with a model selected by ModelFinder and 1,000 ultrafast bootstrap replicates (52). The phylogenetic tree was visualized using ggtree (53).

### Detection of Homing Endonucleases

For group I introns longer than 500 bp, open reading frames (ORFs) were predicted using ORFfinder (https://www.ncbi.nlm.nih.gov/orffinder/). The resulting ORFs were analyzed with hmmscan from HMMER v3.3.2 (http://hmmer.org/) against the Pfam-A.hmm v37.0 (54) (E-value < 0.01) to detect conserved homing endonuclease domains.

### Subgroup Classification of Group I Introns

Subgroups classification was performed following the approach described in a previous study (35), using Infernal (cmscan with the options --max and --tblout). Covariance models for group I intron subgroups were downloaded from https://github.com/LaraSellesVidal/Group1IntronDatabase. The covariance model that yielded the highest alignment score was used to assign the subgroup for each intron.

### Construction of Phylogenetic Tree for Group I introns

Group I intron sequences inserted into tRNA genes were aligned using MAFFT v7.525 (55) with the --auto option, and the resulting alignment was trimmed using the -gappyout option in trimAl (56). Maximum-likelihood phylogenetic trees were then constructed using IQ-TREE v2.1.4-beta with a model selected by ModelFinder and 1,000 ultrafast bootstrap replicates. The phylogenetic tree was visualized using iTOL (57).

## ACKNOWLEDGMENTS

This study was supported by JST CREST JPMJCR20S4, SPRING JPMJSP2108, JSPS KAKENHI 24K00749, and S.S. was partly supported by JSPS KAKENHI 22H05152 and AMED ASPIRE Program 25jf0126009h0002. Computation was supported by the HOKUSAI SailingShip supercomputer facility at RIKEN.

## SUPPLEMENTARY INFORMATION

Supplementary material is available for this article: Supplementary Tables 1-10.

